# Real-time detection of condensin-driven DNA compaction reveals a multistep binding mechanism

**DOI:** 10.1101/149138

**Authors:** Jorine M. Eeftens, Shveta Bisht, Jacob Kerssemakers, Christian H. Haering, Cees Dekker

**Affiliations:** Department of Bionanoscience, Kavli Institute of Nanoscience Delft, Delft University of Technology, Delft, Netherlands; Cell Biology and Biophysics Unit, Structural and Computational Biology Unit, European Molecular Biology Laboratory (EMBL), Heidelberg, Germany

## Abstract

Condensin, a conserved member of the SMC protein family of ring-shaped multi-subunit protein complexes, is essential for structuring and compacting chromosomes. Despite its key role, its molecular mechanism has remained largely unknown. Here, we employ single-molecule magnetic tweezers to measure, in real-time, the compaction of individual DNA molecules by the budding yeast condensin complex. We show that compaction proceeds in large (~200nm) steps, driving DNA molecules into a fully condensed state against forces of up to 2pN. Compaction can be reversed by applying high forces or adding buffer of high ionic strength. While condensin can stably bind DNA in the absence of ATP, ATP hydrolysis by the SMC subunits is required for rendering the association salt-insensitive and for subsequent compaction. Our results indicate that the condensin reaction cycle involves two distinct steps, where condensin first binds DNA through electrostatic interactions before using ATP hydrolysis to encircle the DNA topologically within its ring structure, which initiates DNA compaction. The finding that both binding modes are essential for its DNA compaction activity has important implications for understanding the mechanism of chromosome compaction.

## INTRODUCTION

The Structural Maintenance of Chromosome (SMC) complexes cohesin and condensin play central roles in many aspects of chromosome biology, including the successful segregation of mitotic chromosomes, chromatin compaction, and regulation of gene expression (reviewed in refs 1–3). SMC protein complexes are characterized by their unique ring-like structure (Fig. 1a). The architecture of condensin is formed by a heterodimer of Smc2 and Smc4 subunits, which each fold back onto themselves to form ~45-nm long flexible coiled coils^4^, with an ATPase “head” domain at one end and a globular “hinge” hetero-dimerization domain at the other end^5^. The role of ATP binding and hydrolysis by the head domains has remained largely unclear. The head domains of the Smc2 and Smc4 subunits are connected by a protein of the kleisin family, completing the ring-like structure (Fig. 1a). The condensin kleisin subunit furthermore recruits two additional subunits that consist mainly of HEAT-repeat motifs. Most metazoan cells express two condensin complexes, condensin I and II, which contain different non-SMC subunits and make distinct contributions to the formation of mitotic chromosomes^6^. The budding yeast *Saccharomyces cerevisiae*, however, has only a single condensin complex, which contains the kleisin subunit Brn1 and the HEAT-repeat subunits Ycg1 and Ycs4 (Fig. 1 a).

**Figure 1:**
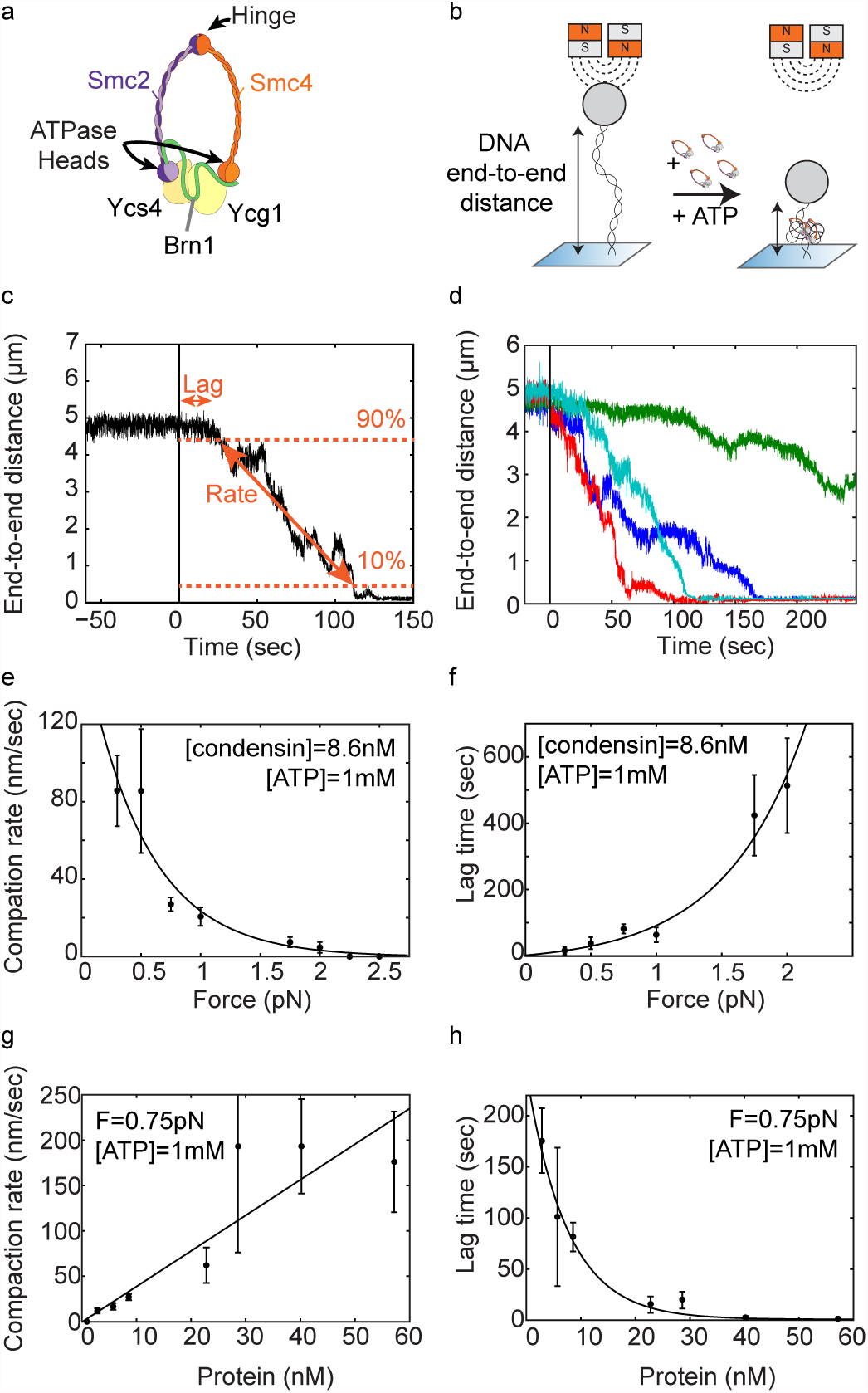
Condensin compacts DNA in the presence of ATP. **a)** Cartoon of the yeast condensin complex. Smc2 and Smc4 dimerize via their hinge domains. The kleisin Brn1 associates with the head domains to create a ring-like structure. HEAT-repeat subunits Ycs4 and Ycg1 bind to Brn1. **b)** Schematic representation of the compaction experiment. A DNA molecule is tethered between a glass slide and a magnetic bead. When condensin and ATP are added, the end-to-end length of the DNA decreases. **c)** Characterization of the compaction process with two parameters. The lag time is defined as the time it takes for the compaction to initiate. The compaction rate is set by the compaction speed between 90% and 10% of the original end-to-end length. **d)** Examples of compaction traces. Each color represents a different individual DNA tether measured in the same experiment. Condensin (8.6nM) and ATP (1 mM) are added at time point zero. **e)** The average compaction rate decreases as force increases. At forces higher than 2pN, condensin does not compact DNA. At 2pN, 2 out of 9 tethers did not condense. At 1.75pN, 2 out of 8 tethers did not condense. Error bars represent SEM. For all these experiments, condensin concentration was 8.6nM and ATP concentration was 1mM ATP. An exponential curve (line) is added as a guide to the eye. **f)** The lag time increases as force increases. An exponential curve (line) is added as a guide to the eye. **g)** The average compaction rate increases linearly as protein concentration increases. Asterisk indicates that not all DNA tethers showed compaction at that protein concentration. At 2.86nM, 5 out of 15 tethers in the experiment did not compact. **h)** The lag time decreases as protein concentration increases. An exponential curve (line) is added as guide to the eye.

How condensin complexes associate with chromosomes has remained incompletely understood. Biochemical experiments have provided evidence that condensin, similar to cohesin, embraces DNA topologically within the ring formed by the Smc2, Smc4 and kleisin subunits^7^. In addition, the HEAT-repeat subunits were found to contribute to condensin’s loading onto chromosomes and the formation of properly structured chromosomes^8^,^9^. In contrast, ATP hydrolysis by the Smc2–Smc4 ATPase heads does not seem to be absolutely required for the association of condensin with chromosomes *in vivo*^10^ and condensin binds DNA *in vitro* even in the absence of ATP^7,11^. DNA, however, can stimulate ATP hydrolysis by the Smc2–Smc4 ATPase heads^8,12^. These findings have led to the speculation that condensin might initially bind to the DNA double helix by a direct interaction with its HEAT-repeat and kleisin subunits and that this binding might subsequently trigger an ATP hydrolysis-dependent transport of DNA into the condensin ring^7^. Such a hypothesis has not yet been confirmed, however. The condensin-DNA interaction is presumably the key to the mechanism by which condensin drives DNA compaction, a subject of keen interest that is intensely debated (reviewed in refs 13,14). Models for the condensin-driven compaction of DNA include random crosslinking, condensin multimerization, and/or DNA loop extrusion^15–20^. The loop extrusion model has recently gained support, but a consensus has not yet been reached^21^. Finally, condensin has also been suggested to alter the supercoiled state of DNA to promote DNA compaction ^22–25^.

One caveat of most biochemical experiments is that they can only probe the final geometry of the DNA, but cannot address the interaction of condensin molecules with DNA during the compaction cycle. To truly resolve the compaction mechanism, an understanding of the binding properties of individual condensin complexes to DNA will be essential. Single-molecule techniques are especially suitable for investigating the mechanical properties, structure, and molecular mechanism of SMC proteins. For example, single-molecule imaging methods proved to be crucial for revealing the sliding and motor action of individual SMC complexes on DNA^26^^-^^29^. Likewise,magnetic tweezers experiments have been successfully used to describe the compaction of DNA by the *Escherichia coli* SMC protein MukB^30^ and by condensin I complexes immunoprecipitated from mitotic Xenopus laevis egg extracts^11^.

To obtain insights into the DNA compaction mechanism of condensin complexes, we here employ magnetic tweezers to study DNA compaction induced by the *S. cerevisiae* condensin holocomplex at the single-molecule level. Magnetic tweezers are exquisitely fit to study the end-to-end length and supercoiling state of DNA at the single-molecule level. We show real-time compaction of DNA molecules upon addition of condensin and ATP. The compaction rate depends on the applied force and the availability of protein and hydrolysable ATP. Through rigorous systematic testing of experimental conditions, we show that condensin makes a direct electrostatic interaction with DNA that is ATP independent, while only upon ATP hydrolysis, condensin does associate with DNA in a salt-resistant, most likely topological binding mode, where the DNA is fully encircled by the condensin ring. Our findings are inconsistent with a “pseudo-topological” binding mode, in which a DNA molecule is sharply bent and pushed through the condensin ring without the need to open the SMC–kleisin ring. Our results show that condensin uses its two DNA-binding modes to successfully compact the DNA, thus setting clear boundary conditions that must be considered in any DNA organization model. We present a critical discussion of the implications of our results on the various models for the mechanics of condensin-mediated DNA compaction and conclude that our findings are consistent with a loop-extrusion model.

## RESULTS

### Condensin compacts DNA molecules against low forces

To measure the real-time compaction of individual linear DNA molecules by the *S. cerevisiae* condensin holocomplex in a magnetic tweezers set-up, we tethered a DNA molecule between a magnetic bead and a glass surface in physiological buffer conditions (Fig. 1 b). We then used a pair of magnets to stretch the DNA and apply force. We routinely performed a pre-measurement to determine the end-to-end length of the bare DNA at the force applied (Fig. 1c, left of the black vertical line at time point zero). We then simultaneously added condensin (8.6nM) and ATP (1 mM) to the flow cell (time point zero, Fig. 1c). Following a short lag time, the end-to-end length of the DNA started to decrease until, in the vast majority of cases, the bead had moved all the way to the surface. We thus observe condensin-driven DNA compaction in real-time at the single-molecule level.

We quantitatively characterized the condensation process using two parameters. First, we measured the lag time: the time it takes for compaction to initiate after adding condensin at time zero (Fig. 1c). Second, from the decrease in the end-to-end length of the DNA, we measured the compaction rate in nanometers per second (Fig. 1c). To avoid a bias at either end of the curve, we extracted the average compaction rate from the decrease between the 90% and 10% levels of the initial end-to-end length. Different DNA tethers in the same experiment typically displayed a sizeable variation between individual compaction traces (Fig. 1d).

While keeping protein and ATP concentrations constant, we measured compaction rates at different applied forces. We found that condensin was able to compact DNA against applied forces of up to 2pN, albeit with rates that strongly decreased with force (Fig. 1e). Note that 2pN is a rather low force for many biological motor proteins. On average, the rate was constant over the course of the experiment for each tether, only slowing down slightly towards the end (Supplementary Fig. S1). Lag times increased with increasing force (Fig. 1f). We conclude that compaction is slower and takes longer to initiate when condensin complexes are acting against a higher applied force.

The compaction rate increased approximately linearly with protein concentration (Fig. 1g). Higher amounts of protein were able to condense DNA much faster, at rates of up to 200nm/sec. Similarly, the lag times decreased at higher protein concentrations (Fig. 1h). Thus, at higher concentrations, multiple condensin complexes might work in parallel on the same DNA molecule, resulting in faster compaction.

### DNA compaction depends on condensin binding and ATP hydrolysis

We found that the compaction rate increased with increasing ATP concentrations and saturated at concentrations above a few mM (Fig. 2a). A Michaelis-Menten fit to the data resulted in a v_max_ of 85 ± 28nm/s (95% confidence interval) and a K_M_ of 1.4 ± 1.5mM. Lag times were much longer at lower ATP concentrations (Fig. 2b).

**Figure 2:**
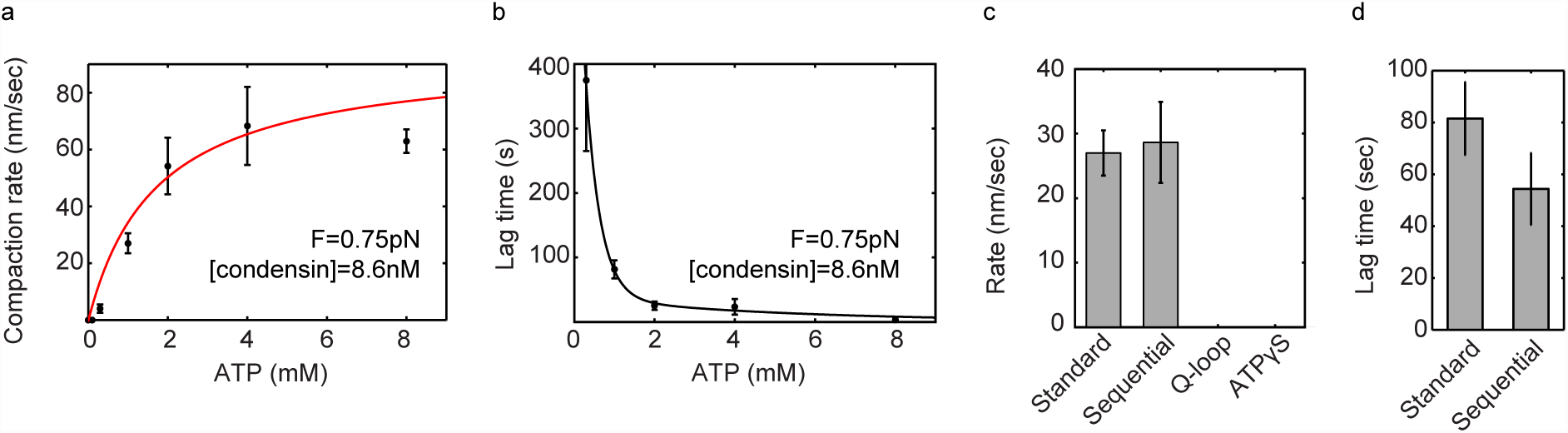
Compaction depends on ATP hydrolysis. **a)** Average compaction rate increases according to a Michaelis-Menten relation with ATP concentration. For this protein concentration, the rate saturates at ATP concentrations higher than 2mM. **b)** Lag time decreases as ATP concentration increases. Line is added as guide to the eye. **c)** Average compaction rate for the standard experiment (0.75pN, 8.6nM protein, 1 mM ATP) and the sequential addition experiment (first 8.6nM protein/no ATP, wash with buffer, and only then add 1 mM ATP). Rate is similar for standard and sequential addition. The ATPase mutant and wild-type in the presence of ATPyS do not show compaction. **d)** The lag time for the standard experiment and the sequential addition experiment. The lag time decreases for sequential addition.

We then tested whether condensin can bind DNA at all in the absence of ATP. We incubated condensin with DNA substrates for 20 minutes in the absence of ATP and, as expected, observed no DNA compaction during this time period (Supplementary Fig. S2a). We then washed the flow cell with buffer without ATP to remove all unbound condensin, before flowing in buffer containing 1 mM ATP but *no* additional protein (hereafter called “sequential addition”). After ATP addition, we observed robust compaction (Supplementary Fig. S2a, N=11) with a rate that was similar to the rate measured when adding protein and ATP simultaneously (Fig. 2c). Interestingly, the lag time was shorter for the sequential addition (Fig. 2d). These experiments are consistent with the notion that condensin can bind in the absence of ATP, remains attached during washing steps, and can start DNA compaction when ATP is subsequently added^11^.

To confirm that compaction needs ATP hydrolysis and not merely ATP binding, we investigated whether condensin would induce compaction in the presence of ATPγS, an only slowly hydrolysable ATP analog, or when ATP hydrolysis was prevented by mutations in the Q-loop motifs of Smc2 and Smc4 (Smc2-Q147L/Smc4-Q302L). As expected, either replacement of ATP by ATPγS or mutation of the ATPase sites abolished DNA compaction in our assay (Fig. 2c and Supplementary Fig. S3). We conclude that both ATP binding and ATP hydrolysis, are essential for the DNA compaction activity that we observe.

### Condensin remains bound to DNA after force-induced de-compaction

We next tested whether the condensin-DNA interaction can be disrupted by applying a high force once the compaction reaction had taken place. Initially, we quantified the end-to-end extension of the bare DNA at 10pN and 0.75pN forces (Fig. 3a). After adding condensin and ATP we observed compaction, as before (Fig. 3b). As soon as the DNA molecule had been compacted to about half of its original length, we abruptly increased the force to 10pN (Fig. 3c). Upon this sudden force increase, the end-to-end length did not immediately recover to the fully extended level, in contrast to the response of a bare DNA molecule. Instead, it took a few seconds (here ~5sec) until the DNA had extended all the way to the end-to-end length we had measured for the bare DNA at 10pN (Fig. 3a). When we subsequently lowered the force to 0.75pN, the DNA started to compact again from the same level it had started at the beginning of the experiment (Fig. 3d). We conclude that condensin-dependent DNA compaction can be fully reversed by stretching the DNA with high forces, as reported previously^11^. This, however, does not hinder subsequent compaction at low force.

**Figure 3:**
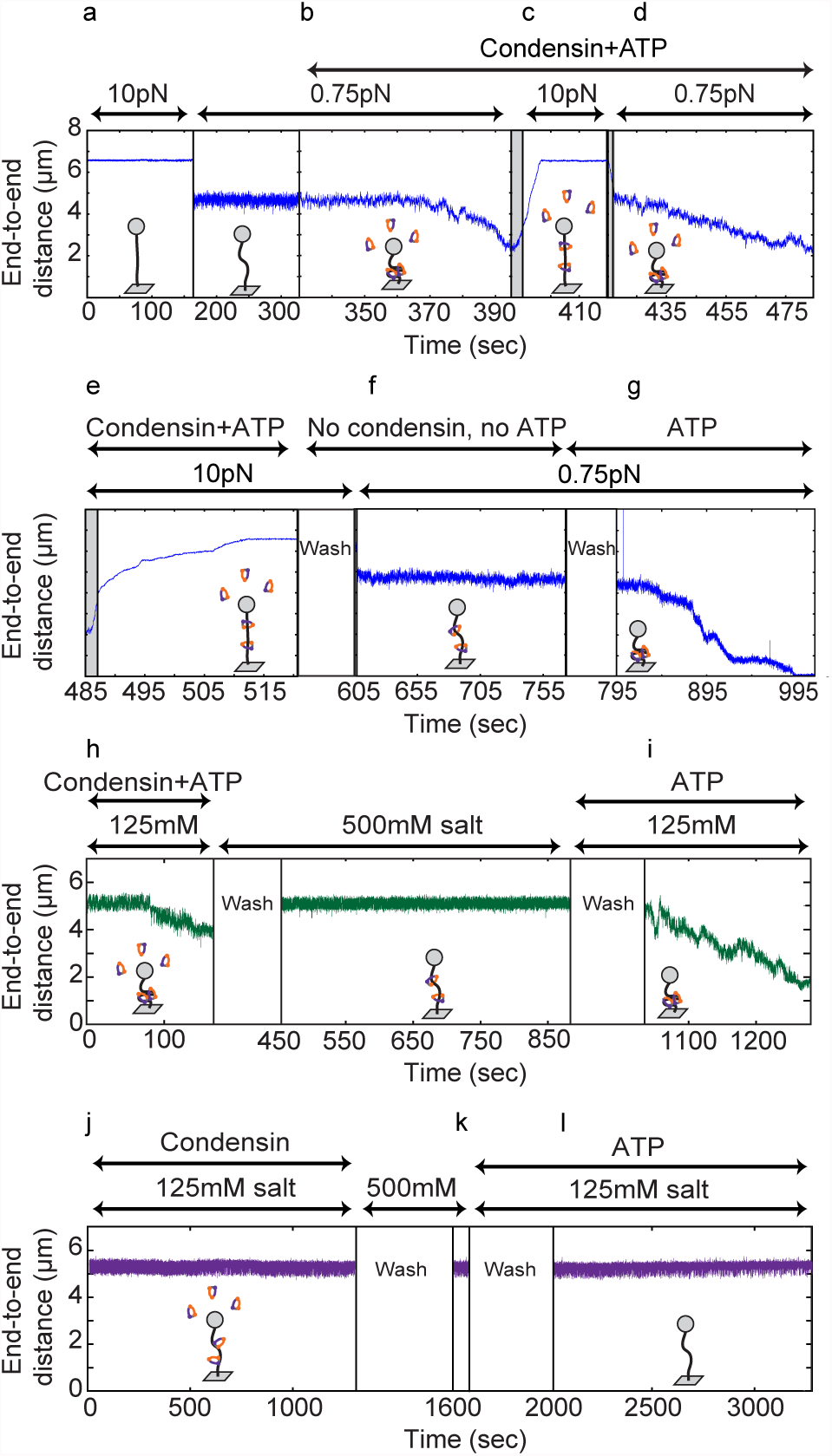
De-compaction by switching to high force or high salt. **a)** Pre-measurements of the DNA tether length before addition of condensin. The average end-to-length was recorded at 10pN (left) and 0.75pN (right). This trace is an example taken from N=28 independent experiments. Although different tethers showed different rates of compaction and de-compaction, the qualitative result was identical for all. **b)** With the force at 0.75pN, condensin (8.6nM) and ATP (1 mM) were added and the DNA was compacted. **c)** After about 50% compaction, the force was suddenly increased to 10pN. The grey area indicates where the time that the magnet was moving and the black vertical line indicates the point the magnet arrived at the 10pN position. DNA end-to-end length increased, reversing compaction and eventually recovering the full end-to-end length. **d)** The force was subsequently lowered to 0.75pN again. Condensin and ATP were still present and the DNA condensed again. **e)** The force was increased to 10pN again. The DNA end-to-end length again increased and eventually recovered to the premeasured full length of bare DNA at 10 pN. Next, the flow cell was washed with buffer without any ATP or protein. **f)** The force was then lowered to 0.75pN, and the DNA was observed to not compact in the absence of ATP. Next, the flow cell was washed with buffer with 1 mM ATP but no protein. **g)** After thus adding ATP but no extra protein, the DNA was able to condense again, indicating that the protein remained bound after pulling and washing. **h)** The green trace shows a different experiment. At time=0s, 8.6nM condensin and 1 mM ATP were added as normal. After compaction, the flow cell was washed with high salt (500mM), and the compacted structure was extended again. **i)** At time=900s, the flow cell was washed with physiological salt and 1 mM ATP but *no* additional protein, and the DNA compacted again. **j)** The purple trace shows a different experiment. At time=0s, 8.6nM condensin but no ATP was added, and no compaction was observed. **k)** The flow cell was then washed with high salt (500mM), and no change in end-to-end length was detected. **l)** The flow cell was washed with physiological salt and 1mM ATP was added. No compaction was observed.

We then repeated the 10pN pulling step, and this time it took even longer (~25 sec) until the DNA recovered the full end-to-end length (Fig. 3e). While keeping the force at 10pN, we then washed the flow cell with buffer *without* ATP or protein to remove all ATP and unbound condensin. When we then lowered the force to 0.75pN, the DNA did *not* compact (Fig. 3f). Strikingly, however, as soon as we added ATP (but *no* additional protein), we again observed compaction (Fig. 3g). This result demonstrates that, first, condensin can stay bound to DNA even when stretching the DNA at high forces and washing with physiological buffers and, second, that condensin that had remained bound to DNA requires ATP to initiate a new round of DNA compaction. We confirmed the findings such as those outlined in Figure 3a-g in many independent experiments (N=28).

### Condensin associates with DNA in two distinct binding modes

Condensin might mediate DNA compaction through direct electrostatic interactions with the DNA helix, through topologically encircling DNA within its ring structure, through pseudo-topologically entrapping DNA by inserting a DNA loop into its ring (Supplementary Fig. S6c), or through a combination of these modes. Whereas electrostatic interactions are sensitive to high salt concentrations, bulk biochemistry experiments have shown that condensin’s topological interaction is resistant to salt concentrations of 500-1000mM NaCl^7^. We therefore assayed whether compaction remained stable after washing with buffer containing 500mM NaCl after compaction had been achieved by condensin and ATP. We found that DNA compaction was fully reversed by the high salt conditions (Fig. 3h, t=450sec, N=7). This indicates that electrostatic interactions with DNA are required for maintaining the condensin-mediated compaction state of DNA. Strikingly, when we subsequently lowered salt concentrations to physiological levels (125mM NaCl) in the presence of ATP (but without additional protein), we again observed compaction (Fig. 3i, t=1050sec). This demonstrates that condensin, once it had been loaded onto DNA by use of ATP, remained associated with the DNA during the high salt wash and was capable of again compacting DNA in an ATP-dependent reaction once salt concentrations had been lowered.

We next tested whether ATP was required to allow condensin to bind DNA in a salt-resistant manner. We first incubated condensin with DNA in physiological buffer conditions without ATP (as in the sequential addition experiment). As expected, we observed no compaction in the absence of ATP (Fig. 3j, t=0-1300sec). We then washed with high salt buffer (500mM NaCl) before lowering salt concentrations again to 125mM and adding ATP (Fig. 3k). In contrast to the previous experiment where condensin had been allowed to bind DNA in the presence of ATP before the high salt wash, we did not observe any compaction (Fig. 3l, N=9). Similarly, when we incubated condensin with DNA in the presence of ATPγS instead of ATP, we did not observe compaction (N=14). These experiments demonstrate that ATP hydrolysis is required to convert condensin from a salt-sensitive to a salt-resistant binding mode, which is indicative of topological binding.

We finally examined whether continued ATP hydrolysis was necessary to maintain DNA in the compacted form, since it had been reported that continuous ATP hydrolysis is necessary to maintain the structure of mitotic chromosomes^9^. When we interrupted ongoing DNA compaction by flushing with buffer without ATP, compaction did neither continue nor reverse. Instead, the DNA end-to-end length remained stable (Supplementary Fig. S2b, N=9). When we added ATP again, compaction proceeded. These data demonstrate that the presence of ATP is required to initiate and continue compaction, but is neither necessary for maintaining condensin’s association with DNA nor essential for preserving already compacted DNA structures.

### Condensin compacts DNA in a stepwise manner

Many compaction traces showed sudden distinct decreases in the DNA end-to-end length, to which we will refer as steps. We used a well-established user-bias-independent step-finding algorithm to extract the size of these compaction steps (see Methods and SI for details). In brief, this algorithm objectively evaluates if a trace shows steps, without prior knowledge of step size or location, based on chi-squared minimization. Figure 4a shows a typical example of a DNA compaction trace with fitted steps. We used this hands-off algorithm to analyze all traces we had collected and determine the step sizes.

**Figure 4:**
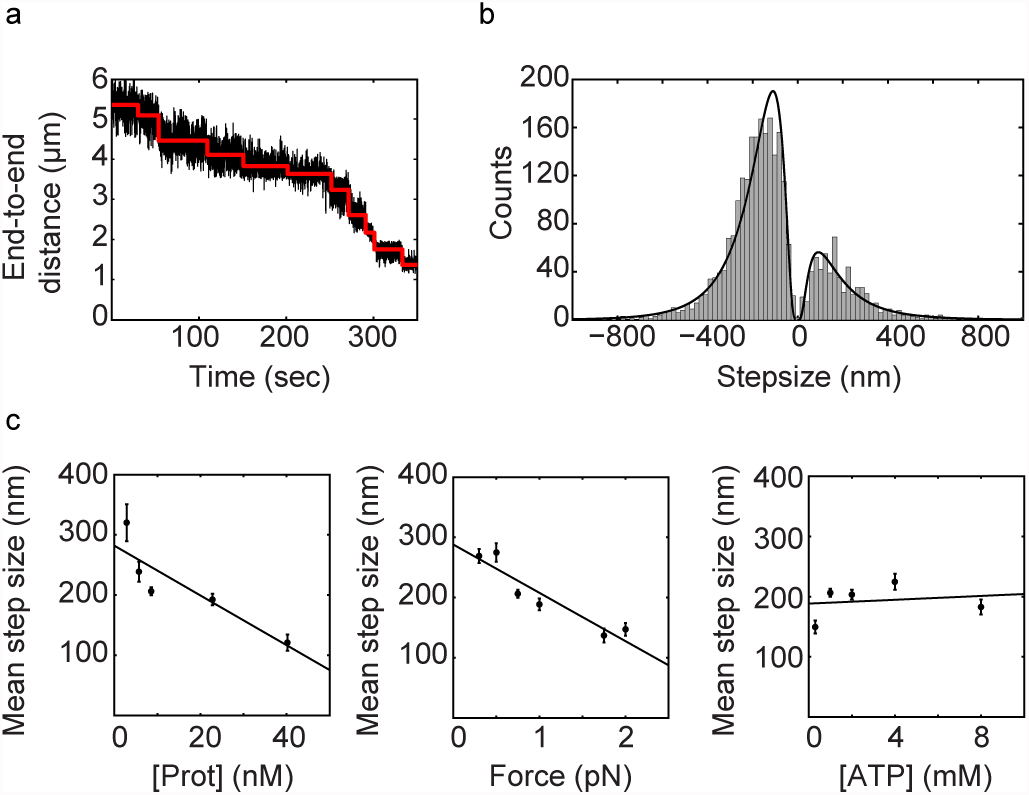
Condensin compacts DNA in a stepwise manner. **a)** Compaction occurs with discrete steps. Black trace depicts the raw data; red is the fitted step trace. **b)** Histogram of compaction and de-compaction steps, measured from all traces included in this paper. Lognormal distributions are fitted. Negative peak: mean= −205nm, SD=148nm, SEM=3nm. Positive peak: mean=213nm, SD=185nm, SEM=7nm. **c)** Step size decreases with protein concentration and force. ATP concentration does not affect the step size. Linear fits are added as guide to the eye. Error bars represent S.D. For this figure, only negative steps were considered.

Figure 4b shows a histogram of the step-size distribution (2675 steps from 90 traces). The size distribution histogram displays two peaks, corresponding to compaction steps (negative step values, N=2045) and less abundant spontaneous de-compaction steps (positive step values, N=630). The data is well described by a lognormal distribution, with mean values of 205 ± 148nm (negative steps, mean ± SD) and 213 ± 185nm (positive steps, mean ± SD). Note that these are remarkably high values, which are clearly larger than the size of a condensin molecule itself (see Discussion). The distributions are wide, indicating that there is a range of outcomes for individual compaction steps. Notably, traces resulting from force-induced de-compaction (10pN, Fig. 3) were smooth and did not show any discernable steps (and were accordingly rejected by our step-finding algorithm). We generated step size histograms for all conditions and fitted lognormal distributions (Supplementary Fig. S4), which showed that the mean step size decreased linearly with protein concentration and with force. In contrast, the step size did not depend on the ATP concentration.

### Condensin does not compact DNA by inducing DNA supercoiling

As condensin was reported to influence the supercoiled state of plasmid DNA in the presence of topoisomerases^22–25^, it has been proposed that condensin might actively introduce (positive) supercoiling into DNA helices to promote their compaction. We therefore examined the compaction activity as a function of the DNA supercoiling state, an assay for which magnetic tweezers are especially suitable. An example of a rotation curve for a torsionally constrained DNA molecule is shown in Figure 5a. On average, half of the DNA tethers in each experiment were torsionally constrained (and hence can be used to test possible effects of supercoiling on the compaction), while the other half did not show a decrease in end-to-end length upon rotation due to a nicked tether. Remarkably, we found no differences in the compaction rates between nicked and torsionally constrained DNA molecules (Fig. 5b). We also tested whether the initial topological state of the DNA would affect the compaction process by introducing +30 or –30 turns into the DNA molecules before adding protein and ATP. Again, we did not find a measurable effect on the compaction rate (Fig. 5b). If the decrease in the end-to-end length during compaction were due to condensin introducing supercoils, we should be able to actually *extend* DNA that previously was compacted by condensin, as condensin would remove some of the applied supercoils^31^. We therefore applied +50 or –50 turns to a DNA molecule that was halfway compacted (Supplementary Fig. S5a). Upon starting the rotation curve to either direction, we never observed that the DNA end-to-end length increased, but instead measured a decrease in compaction in both cases (Supplementary Fig. S5b).

**Figure 5:**
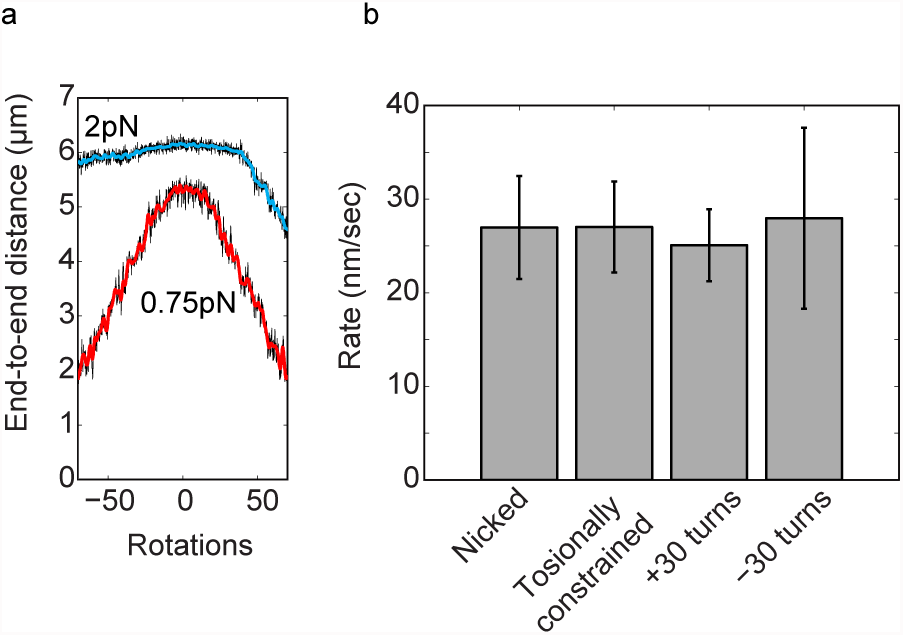
DNA supercoiling does not influence compaction. **a)** Rotation curves of a bare DNA molecule at constant forces (2pN in blue, 0.75pN in red), showing that this molecule is torsionally constrained and supercoils are introduced by applying positive or negative rotations to the magnets. **b)** Compaction rates for different supercoiling states. All measurements are for the standard experiment: 0.75pN, 1 mM ATP, 8.6nM condensin. There is no difference between nicked DNA and coilable DNA. Also there is no difference between relaxed DNA, and DNA with applied turns in either direction.

These findings show that the condensin-induced decrease in compaction was not a result of DNA supercoiling. However, when we rotated the magnet back to the starting position (0 turns) after applying 50 turns to compacted tethers, we found that the end-to-end length did not fully recover. In fact, the end-to-end length started to decrease further already before the “relaxed” point at 0 turns. This behavior occurred regardless of the direction in which the DNA had initially been rotated (N=8, both directions). We speculate that instead of actively introducing supercoils, condensin is able to “lock” DNA plectonemes by embracing their stem (see Supplementary Fig. S5c).

## DISCUSSION

### DNA binding and compaction are distinct steps in the condensin reaction cycle

We used single-molecule magnetic tweezers to demonstrate that condensin holocomplexes purified from *S. cerevisiae* are able to compact DNA, similar to a previous study of condensin I complexes immunopurified from *X. laevis* egg extracts^11^. In contrast to this previous report, DNA compaction clearly dominated any de-compaction in our assays.

Our data show that association of condensin with DNA can take place in the absence of ATP (Supplementary Fig. S2a). This ATP-independent interaction is able to survive washing steps with physiological salt concentrations, but it does not survive in buffer conditions of high ionic strength (Fig. 3j, k), indicative that this interaction of condensin with DNA might be electrostatic. We propose that this binding step occurs through the direct interaction with the DNA double helix of the condensin HEAT-repeat and kleisin subunits^8^ and/or possibly through another yet unidentified DNA binding site in the complex (Fig. 6). Notably, condensin apparently does not require any loading factor(s) to associate with and to compact DNA. This contrasts cohesin, which commonly uses specific loading factors to increase the efficiency of its binding to chromosomes^26,27,32,33^.

**Figure 6:**
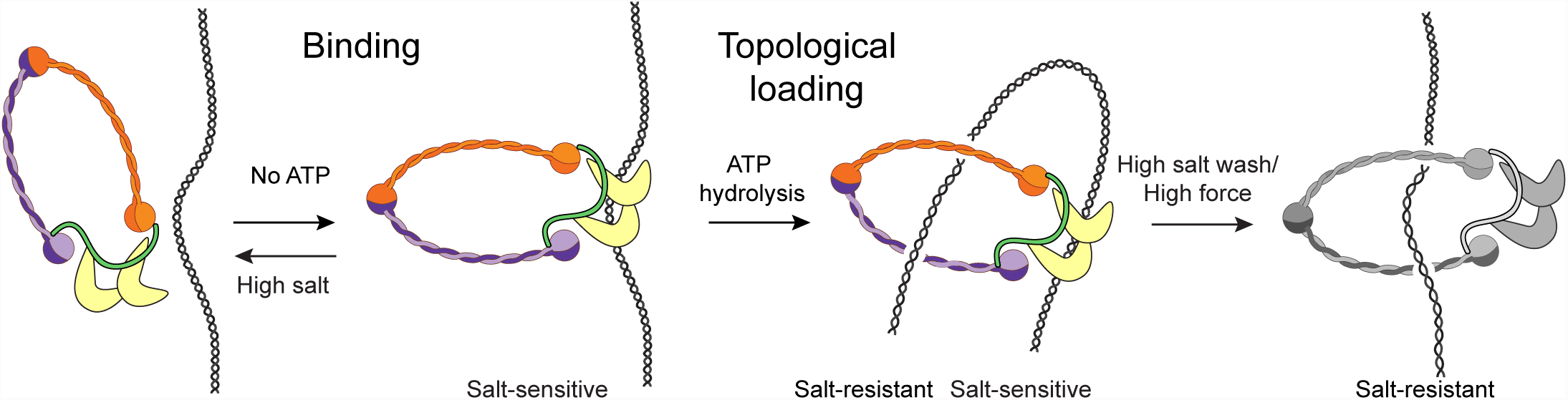
Condensin compacts DNA using a multistep binding mechanism. We propose a multistep model for condensin’s binding mechanism, represented by the colored panels. First, condensin binds to DNA electrostatically, presumably through the HEAT-repeat subunits. Next, upon ATP hydrolysis, condensin embraces the DNA topologically. This initiates the compaction of DNA. Finally (most right panel in gray), in our experiments, we disrupted the electrostatic interactions in some of our assays with high salt or high force.

When condensin is added to DNA in the presence of ATP, however, it is able to survive high salt conditions (Fig. 3h, i). This suggests that the ATP-dependent mode of DNA binding must be exceptionally stable, e.g. such as provided by a topological binding mode where condensin encircles the DNA. In contrast to the initial step of DNA binding, the subsequent compaction step essentially depends on ATP hydrolysis by the Smc2–Smc4 subunits of the condensin complex. The importance of ATP hydrolysis, not just ATP binding, is evident from the deficiency of the ATPase mutant to compact the tethered DNA substrates in our assay. It thus appears logical to conclude that the electrostatic interaction is converted into a topological interaction by ATP-dependent temporary ring opening and entry of the bound DNA into the ring (Fig. 6). Possibly, the initial electrostatic interaction might release upon ATP hydrolysis and this binding site would then again be available to grab another piece of the same DNA and thereby create a DNA loop.

### ATP hydrolysis by the Smc2–Smc4 subunits is required for stepwise DNA compaction

Following a short lag time after addition of ATP, condensin induces a fast stepwise compaction of the DNA tethers. We interpret the lag time before compaction starts as the time necessary for condensin to become active for compaction, which likely involves the conversion of an electrostatic into a topological binding mode. This is consistent with the finding that the lag time depends on the concentration of ATP, in addition to the protein concentration. The observation of a lag time is furthermore consistent with recent measurements of condensin movements on DNA curtains, where we see condensin bind and pause before becoming active for translocation^28^. Notably, a mutant version of condensin that is unable to hydrolyze ATP displays no compaction activity in our assay. This suggests that the required energy for compaction must stem from ATP hydrolysis by the Smc2–Smc4 subunits. While ATP hydrolysis is essential for the compaction process, we find that condensed DNA remains compacted even after washing with buffer that does not contain ATP. This shows that continuous ATP hydrolysis by condensin is not required to maintain the compact state of the DNA.

The amount of work needed for compaction can be calculated as the product of the displacement against the applied force. Taking into account that kBT=4.1 pN*nm and that the free energy resulting from hydrolysis of one DNA molecule of ATP is 20kBT, we can calculate the amount of ATP molecules that would minimally be required to drive compaction against a certain force. Assuming for the sake of argument that condensin converts the energy from ATP hydrolysis with 100% efficiency, we estimate that full compaction (from 5 to 0μm) against a force of 0.75pN requires the hydrolysis of 46 ATP molecules, or equivalently, a single 200nm step would require energy from 1.8 ATP molecules. As the force increases, more ATP needs to be hydrolyzed to provide the necessary energy in order to achieve compaction. Accordingly, we find that the step size decreases with increasing force. We also find that the step size decreases with increasing protein concentration, indicating that more complexes are apparently not able to make larger steps. This indicates that clustering of multiple condensin complexes is unlikely to be responsible for the observed compaction activity, since we would expect that larger clusters might make larger steps. Note also that a higher protein concentration does result in faster rates, suggesting that multiple proteins are working in parallel at higher protein concentrations.

A surprising finding from our experiments is the very large size of the compaction steps (50-500nm). It is puzzling that the step size can be larger than the size of the condensin complex, which measures about 70 nm along its longest axis5. One explanation for this conundrum might be that condensin could be taking smaller individual compaction steps and that the steps that we are detecting are in fact bursts of smaller steps. For example, a 200nm step could consist of 4 sequential steps of ~50nm that occur with such a high speed that they cannot be resolved within the temporal resolution of our magnetic tweezer assay (0.1-0.4sec, see Supplementary Methods)^34^. An alternative explanation for the observed step sizes might be that we are measuring the concerted action of not a single but several condensin complexes that act on the same DNA tether in sync, but it remains unclear how the action of multiple condensin complexes could be so precisely coordinated.

### Consequences for geometric models for condensin-induced DNA compaction

Which of the various geometric models for condensin’s mechanism are compatible with our findings? *Generation of DNA supercoiling* has been proposed as a mechanism to condense DNA^24^. Our data are not consistent with this model, since the *S. cerevisiae* condensin was unable to induce DNA supercoils in our assay. Instead, our results indicate that condensin might stabilize or ‘lock’ plectonemes, for example by binding specifically to crossed DNA segments at the stem of DNA plectonemes (Supplementary Fig. S5c). However, while such a mechanism would allow condensin to stabilize an already compacted DNA state, it is unable to induce compaction on its own and hence cannot explain the observed compaction activity.

The *random crosslinking* model proposes that condensin compacts DNA by randomly connecting different pieces of the same DNA molecule^35^ (Supplementary Fig. S6a). Such a scenario fits well with a large distribution of step sizes as well as with step sizes that are considerably larger than the dimensions of the condensin complex itself. This model requires, however, that distant DNA regions come into close proximity for cross-linking. Since, at a force of 1 pN, the DNA tethers are already stretched to 85% of their contour lengths, it is difficult to imagine how, under these forces, large loops could be generated through random cross-linking. Furthermore, since this model does not involve a catalytic compaction activity, it does not explain how halfway compacted DNA molecules can compact further after any free protein has been washed away, as it is quite unlikely that this would happen by condensin letting go of one piece of DNA to grab another piece of DNA further away in order to create a larger loop. Theoretical modeling of the biophysics of a crosslinked DNA polymer under an applied force would be helpful to estimate these notions quantitatively. A variation of the random crosslinking model might involve individual condensin complexes that mutually interact to generate a DNA loop, i.e., in a variation of the handcuff-like model that has been proposed for the cohesin complex^36^. Note that individual condensins must still interact topologically with the DNA to explain the observed salt resistance. This model is in fact consistent with the decreasing step size for increasing protein concentrations, as the changes of finding a second condensin molecule close by to handcuff with are larger for higher protein concentrations. Yet, the model also faces the same challenge of explaining how halfway compacted DNA molecules can compact further after any free protein has been washed away.

A model that recently gained much attention is *loop extrusion*. Here, condensin binds to DNA and moves it through its ring to extrude a loop of DNA, which thereby continuously increases in size (Supplementary Fig. S6b)^37^. Simulations have shown that loop extrusion can indeed achieve efficient chromosome condensation^17^. Requirements for this model are that condensin has DNA motor activity, which was demonstrated recently^28^, and that the extrusion machine can interact with at least two points along the DNA simultaneously. If the interaction of condensin with DNA would only be topological, loop extrusion would not work, as DNA can slip out of the ring, which certainly will happen under an applied force. Our finding that a direct (electrostatic) contact between condensin and DNA is required to maintain the compacted state of DNA suggests that such a contact might serve as an anchor site at the base of a forming loop. The finding that halfway compacted DNA molecules can eventually compact fully without the addition of extra protein is furthermore easy to imagine for a motor extruding a loop of ever-larger size.

Cartoons of the loop extrusion mechanism often depict a pseudo-topological embrace of the DNA (Supplementary Fig. S6c). For such pseudo-topological loading, the condensin ring does not necessarily have to open, in contrast to real topological loading. Importantly, we find that a pseudo-topological embrace is inconsistent with our data, as such a conformation would not survive high salt washes and high force. Instead, our data indicate that the ATP-hydrolysis-assisted DNA loading is truly topological. This is an important distinction that changes the way one should think about loop extrusion, and we accordingly suggest that one should take the topologically loaded state as the basis for future modeling of the loop extrusion process (Fig. 6 and Supplementary Fig. S6c).

In conclusion, systematic evaluation of DNA compaction by condensin complexes allowed us to resolve the binding mode conditions that must be met in any geometric model. Our data demonstrate a two-step model: first ATP-independent direct interaction of condensin with DNA, followed by ATP-hydrolysis-dependent topological loading and DNA compaction (Fig. 6). This model provides an important stride forward in unraveling the mechanism of chromosome compaction by condensin complexes.

## METHODS

### Protein purification

Wild-type and ATPase mutant (Smc4(Q302L)–Smc2(Q147L)) versions of the *S. cerevisiae* condensin holocomplex were overexpressed from galactose-inducible promoters in budding yeast. The complexes were purified from whole cell extracts via a tandem affinity chromatography strategy, using a His12 tag fused to the Brn1 subunit and a triple StrepII tag fused to the Smc4 subunit, followed by a gelfiltration step. Expression and purification are described in detail in ref. 28. Purified proteins were aliquoted, snap frozen in liquid nitrogen, and stored at –80°C.

### Magnetic tweezers

We used a multiplexed magnetic tweezers as described in refs 37 and 38. We used a 20kb DNA construct with digoxygenin- and biotin-handles and nitrocellulose coated flow cells (volume 30μlc39. In brief, nitrocellulose-coated flow cells were incubated with 100mM anti-digoxygenin antibodies (Fab-fragment, Roche). Then, the flow cell was washed with washing buffer (20mM TRIS-HCl pH7.4, 5mM EDTA). Next, the surface was passivated with 10mg/ml BSA for 1 hour and washed again. Streptavidin-coated beads (MyOne, Life Technologies) were incubated with biotin-functionalized DNA for 20 minutes. After incubation, the beads were washed three times with washing buffer plus 0.05% Tween. An excess amount of beads with digoxygenin-functionalized DNA was then incubated in the flow cell for 10 minutes. Finally, the flow cell was washed extensively with compaction buffer (10mM HEPES-NaOH pH7.9, 125mM NaCl, 5mM MgCl2, 1 mM DTT) to flush out all unbound beads and provide near-physiological reaction conditions. Compaction was only observed at conditions around physiological salt concentrations (50-250mM NaCl, data not shown). Different forces were applied by linear translation of the magnets, while rotation of the magnets was used to apply supercoils. A force calibration curve was generated to correlate the magnet height to the force. Before all experiments, all tethers were routinely checked for coilability and for their end-to-end length before starting the compaction reaction (pre-measurement).

### Determination of the compaction-rate and lag-time

All compaction experiments were carried out in compaction buffer (10mM HEPES-NaOH pH 7.9, 125mM NaCl, 5mM MgCl2, 1mM DTT). Different concentrations of ATP and of the *S. cerevisiae* condensin holocomplex (nanomolar range) were dissolved in 50μl of compaction buffer and flushed in, which typically took 15 seconds. Tracking of the beads was started immediately after flushing in the protein and the force was kept constant throughout the experiment. The lag-time was defined as the time it took for the compaction to start. The time points at which the DNA reached 90%, 80%, 70%, etc. of their original end-to-end length (taken from the pre-measurement) were automatically recorded by our custom-made software. The compaction rate was determined by calculating the difference in end-to-end length between the 90% and 10% time points. In the case that compaction did not reach the 10% point, we determined the rate from the initial part of the compaction curve. The standard duration of an experiment was 20 minutes.

### Step analysis

We used a well-defined step-fitting algorithm that was previously described^40^. This algorithm objectively evaluates if a trace shows steps, without prior knowledge of step size or location, based on chi-squared minimization. To evaluate the variation of step sizes in an objective manner, we improved the implementation of this algorithm to allow for hands-off, batch style analysis. For details, see Supplementary Methods.

## ACKNOWLEDGEMENTS

We thank Ana Mota for preliminary research, Allard Katan, Jekyung Ryu, and Jakub Wiktor for discussions, and Damien D'Amours for plasmids and yeast strains for overexpression of the condensin holocomplex. This work was supported by the ERC Advanced Grant SynDiv (no. 669598 to C.D.), the ERC Consolidator Grant CondStruct (no. 681365 to C.H.H.), and by the Netherlands Organization for Scientific Research (NWO/OCW) as part of the Frontiers of Nanoscience program. S.B. acknowledges support from an EMBL Interdisciplinary Postdoctoral fellowship (EIPOD) under Marie Curie Actions (COFUND).

## AUTHOR CONTRIBUTIONS

J.E. magnetic tweezers experiments and data analysis. S.B. protein expression and purification. J.E. and J.K. software. J.E., S.B., C.H.H. and C.D. conceived and designed the study and wrote the manuscript.

